# When Gaze-Pattern Similarity May Interfere With Future Memory

**DOI:** 10.1101/2020.06.04.134171

**Authors:** Nathalie klein Selle, Matthias Gamer, Yoni Pertzov

## Abstract

Human brains have a remarkable ability to separate streams of visual input into distinct memory-traces. It is unclear, however, how this ability relates to the way these inputs are explored via unique gaze-patterns. Moreover, it is yet unknown how motivation to forget or remember influences the gaze similarity and memory relationship. In two experiments, we therefore used a modified directed-forgetting paradigm and either showed blurred versions of the encoded scenes (Experiment 1) or pink noise images (Experiment 2) during attempted memory control. Both experiments demonstrated that higher levels of across-stimulus gaze similarity relate to worse future memory. Although this *across-stimulus interference effect* was unaffected by motivation, it depended on the perceptual overlap between stimuli and was more pronounced for different scene comparisons, than scene-pink noise comparisons. Intriguingly, these findings echo the pattern similarity effects from the neuroimaging literature and pinpoint a mechanism that could aid the regulation of unwanted memories.

---

When reminiscing past experiences, old friends, or even a long-gone love, your eyes are likely to wander around. These eye movements do not seem to be chaotic and already 50 years ago it was suggested that our gaze behavior may not only be important for observing the world around us, but also for remembering it (Noton & Stark, 1971). Since then, a considerable amount of research has shown a link between gaze behavior and memory (e.g., Althoff & Cohen, 1999; Heisz & Shore, 2008; Ryan et al., 2000). One interesting phenomenon is that of gaze reinstatement. Several studies have shown that whenever people retrieve a memory of a complex scene picture, they reinstate encoding-related visual exploration patterns (e.g., Foulsham & Underwood, 2008; Underwood et al., 2009). This reinstatement phenomenon seems to be linked to memory performance. Specifically, greater reinstatement of encoding related eye-movements, during recognition testing, has been found to support memory (e.g., Cooper et al., 2017; Foulsham et al., 2012; Foulsham & Kingstone, 2013; Wynn et al., 2019, 2020). Collectively, these studies examined how we visually explore an *identical* scene stimulus during different situations (i.e., encoding and recognition). However, since memory processes do not occur in a vacuum, and usually include memorizing or retrieving a stream of information (e.g., multiple scenes), it is also important to examine how patterns of visual exploration *across* different stimuli relate to subsequent recognition memory.

Although such concept of *across-stimulus similarity* has not yet been examined with respect to visual exploration, there is evidence that the similarity of neural activation patterns relates to memory performance. For example, lower similarity of encoding-related activation patterns in the hippocampus across stimuli (i.e., pattern separation), has been associated with better memory, whereas this relationship was reversed for other medial temporal lobe regions (LaRocque et al., 2013). These findings were expanded upon by demonstrating that reduced pattern similarity within the hippocampus arises from prior learning and benefits subsequent learning by preventing interference between similar memories (Chanales et al., 2017; Favila et al., 2016; Hulbert & Norman, 2015). Clearly, neural and eye-gaze reinstatement are two different, and separately observed, phenomena. Nevertheless, a very recent study has shown that they are positively correlated, with higher levels of gaze reinstatement during mental imagery being accompanied by higher levels of neural reinstatement in visual processing regions (Bone et al., 2018). Such reinstatement of activation patterns in sensory regions is in turn suggested to be coordinated by the hippocampus (Bosch et al., 2014; Pacheco Estefan et al., 2019). Notably, the hippocampus is not only functionally associated with measures of visual exploration (see also Liu et al., 2017; Ringo et al., 1994; Ryals et al., 2015), but also anatomically connected to the oculomotor system (Shen et al., 2016). Consequently, considering that higher across-stimulus *hippocampal-pattern similarity* at encoding predicts worse memory recollection (LaRocque et al., 2013), it raises the question whether across-stimulus *gaze-pattern similarity* shares a similar link with memory.

Further insight into the aforementioned question can be gained from studies assessing the memory-related effects of visual distraction (e.g., Della Sala et al., 1999; Hecht et al., 2016; Quinn & McConnell, 2006). These studies found that passive viewing of a visual distracter disrupts the memory of a previously encoded image, especially when the distracter is more complex and perceptually similar to the encoded image (Borst et al., 2012; Burin et al., 2007). Comparable results were reported with similar versus dissimilar masks (Blalock, 2013). These findings suggest that there needs to be some perceptual overlap – similarity – between the encoded and the distracting stimuli, in order to interfere with memory (see also Clapp et al., 2010; Dolcos et al., 2007; Tremblay et al., 2005). None of the above studies, however, obtained eye-movement measures. Hence, it remains to be determined whether the findings from these visual distraction studies are also related to differences in eye-gaze similarity – specifically, whether more similar distracters induced more similar eye movements and, consequently, might have been more detrimental to memory.

Considering that our eye movements can be actively controlled, the desire to either forget or remember a stimulus may influence the amount of gaze reinstatement. The last two decades have seen a rising interest in *motivated memory* and revealed initial evidence for the intriguing idea that motivational states can shape the nature of our memory representations. Specifically, several studies showed that we can actively suppress memories we wish to forget (e.g., Anderson & Green, 2001; Depue et al., 2007; Fawcett & Taylor, 2008; Wylie et al., 2008), however contradictory evidence exists (Bulevich et al., 2006; Gao et al., 2016; Zwissler et al., 2015). Additional experiments have also attempted to identify the neural predictors and consequences of successful memory control (Anderson & Hanslmayr, 2014; Hu et al., 2015; Rosenfeld et al., 2017; Ward & Rosenfeld, 2017). An interesting, yet untested, hypothesis that can be derived from this literature is that the effects of motivational states on memory are mediated by the control of eye movements. Thus, it is possible that the motivation to either forget or remember a stimulus might influence gaze similarity and, consequently, the hypothesized similarity-memory relationship.

Taken together, the present study was designed to achieve two major goals: (1) To examine whether across-stimulus gaze similarity is related to future memory success. Based on previous fMRI work and visual distraction studies, more similar exploration of different stimuli (during encoding) is predicted to interfere with later memory; (2) To examine whether motivated memory influences the hypothesized similarity-memory relationship. Thus, the obtained knowledge may not only advance our understanding of the relationship between gaze similarity and memory, but may also reveal whether this relation is susceptible to extrinsic motivation. To address these questions, we designed two experiments that relied on a modified directed forgetting paradigm, each entailing two experimental sessions. In short, on each trial of session 1, participants were presented with a scene image (memory-encoding phase), which was followed by a blank screen with a cue (i.e., white, green, or purple circle) instructing them to either do nothing, remember, or forget the previous scene. Immediately following the cue, participants were presented with either a blurred black-and-white version of the previous scene (in Experiment 1) or a pink noise image (in Experiment 2; memory-regulation phase). Two or three days later, in session 2, participants underwent a recognition memory test (see Method below and Fig. 1). In addition to tracking gaze position, skin conductance was measured throughout both experiments^1^, as a measure of stimulus arousal during encoding (e.g., Bradley et al., 2001), mental effort during attempted memory regulation (e.g., Dawson et al., 2016), and recognition memory during testing (e.g., Meijer et al., 2014).

**Fig 1.**
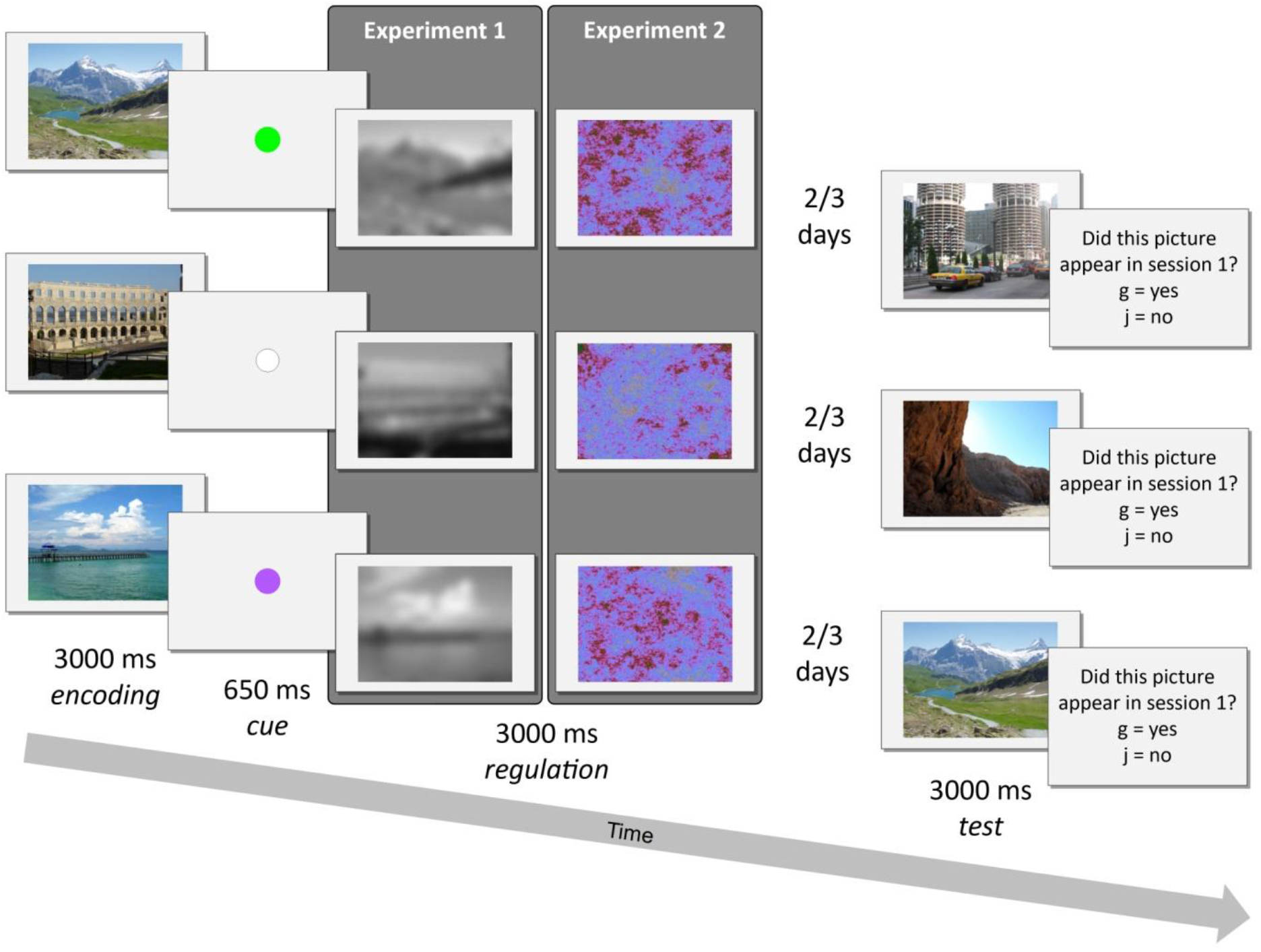
Sequence of events during a remember (green cue), control (white cue), and forget (purple cue) trial of Experiment 1 and Experiment 2. Both experiments entailed two experimental sessions which were separated by two or three days: each trial of session 1 consisted of a memory-encoding and a memory-regulation phase, while session 2 consisted of a recognition-memory test.

## Results

### Experiment 1

#### Recognition performance

Memory accuracy in the second session (i.e., recognition test) was analyzed using a repeated measures ANOVA, comparing recognition rates in the three motivational conditions: remember (*M* = 69%, 95% CI = [63, 75]), control (*M* = 62%, 95% CI = [56, 68]), and forget (*M* = 66%, 95% CI = [60, 71]). The ANOVA yielded a significant effect of motivation, *F*(2,70) = 5.13, *f* = .38 (95% CI = [.10, .62]), *p* = .008, BF_Inclusion_ = 4.9, which was followed by post hoc comparisons. These comparisons revealed significantly higher memory in the remember compared to in the control condition, *t*(35) = 3.49, *p* = .004, *d* = .58 (95% CI = [.11, 1.05]), BF_10_ = 24.3. The other two comparisons (i.e., remember versus forget & control versus forget) did not reach statistical significance (both *p*’s > .05). These results suggest that participants succeeded to enhance (remember > control), but not suppress (forget < control), their memory.

Additional analyses of image memorability revealed that participants forgot different pictures (see online Supplementary Information)^2^, thus indicating that any observed differences in gaze behavior between remembered (hits) and forgotten (misses) images are unlikely to be explained by actual differences between the images themselves.

#### Gaze-similarity analyses

Gaze-pattern similarity was computed using the ScanMatch toolbox for Matlab (The MathWorks, Natick, MA; Cristino et al., 2010). Using this method, each fixation sequence is spatially (12 x 8 bin ROI grid) and temporally (50 ms) binned. Pairs of these sequences are then compared using the Needleman-Wunsch algorithm and the correspondence between pairs is expressed by a normalized similarity score (0 = *no correspondence*, 1 = *identical*; see Fig. 2). Mean similarity scores were analyzed with a Motivation (remember, control, forget) x Memory Accuracy (hits vs. misses) repeated measures ANOVA. Results are depicted in Fig. 3B.

**Fig 2.**
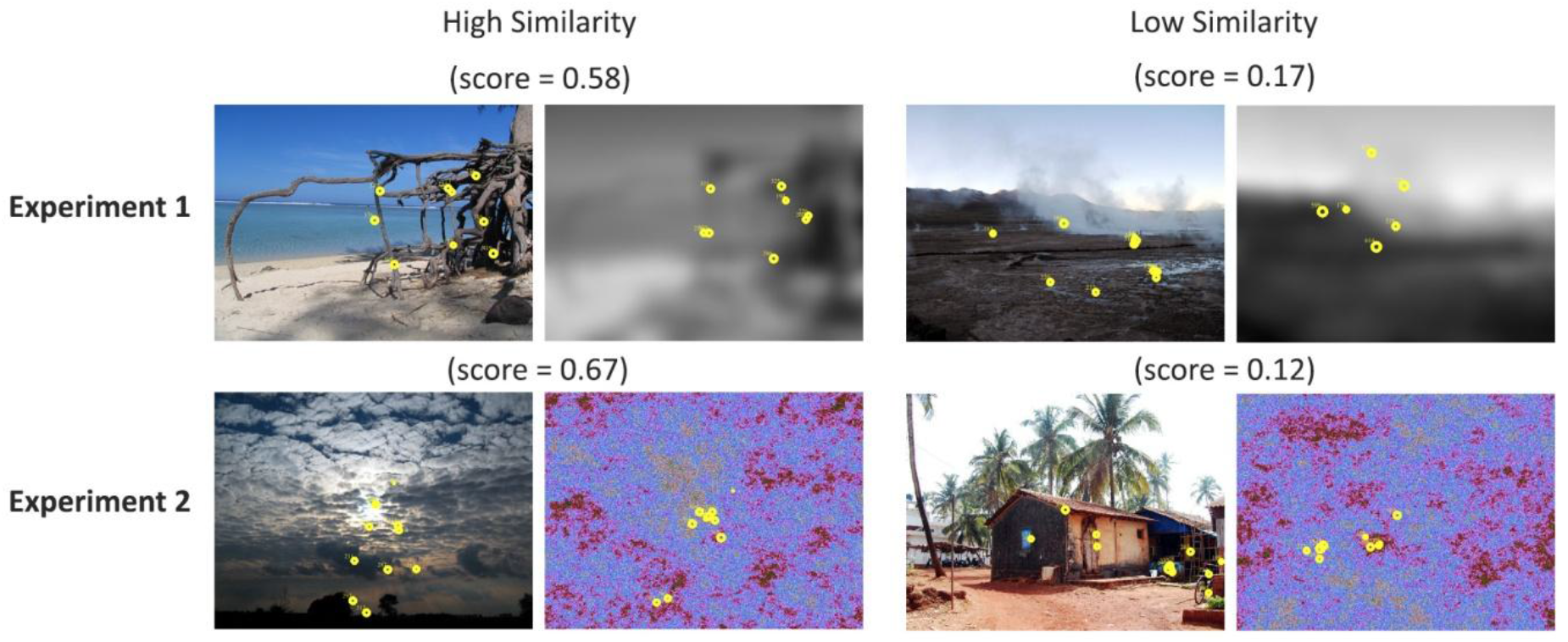
Examples of high and low similarity scores, for both Experiments 1 and 2; each yellow circle is a fixation.

**Fig 3.**
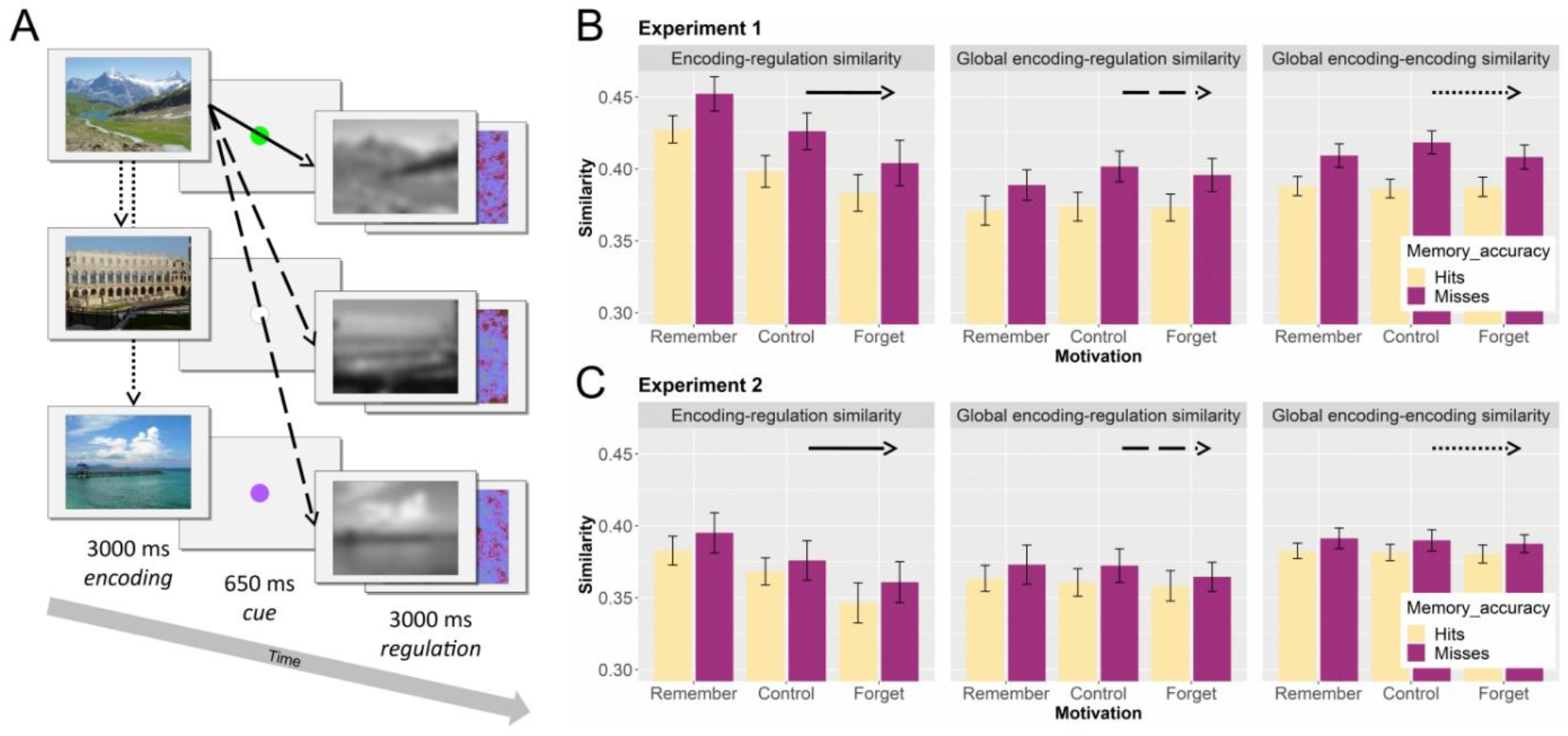
Experimental procedure and results of three across-stimulus similarity measures. Panel A: Sequence of events during session 1 of Experiments 1 and 2. The black arrows illustrate the relevant comparisons for 3 similarity types: solid arrow shows encoding-regulation similarity, dashed arrows show global encoding-regulation similarity, and dotted arrows show global encoding-encoding similarity; Panel B: Bar-plots showing the results of the three similarity measures for Experiment 1; Panel C: Bar-plots showing the results of the three similarity measures for Experiment 2. Means and standard errors (errors bars) across subjects are presented separately for each level of memory accuracy and motivation.

##### Encoding-test similarity

Encoding-test similarity was assessed by comparing the gaze pattern of each image in the memory-encoding phase to the gaze pattern of the same image in the memory-test phase. The ANOVA revealed a significant main effect of motivation, *F*(2,66) = 4.30, *f* = .36 (95% CI = [.06, .61]), ε = .83, *p* = .025, BF_Inclusion_ = 1.4 (which is inconclusive). Post hoc comparisons revealed that the similarity scores in the control condition were significantly higher than the similarity scores in the forget condition, *t*(33) = 3.47, *p* = .004, *d* = .60 (95% CI = [.11, 1.08]), BF_10_ = 22.8. No difference was found between the remember and forget nor between the remember and control conditions (both *p*’s > .05). In addition, the ANOVA revealed a significant main effect of memory accuracy, *F*(1,33) = 9.15, *f* = .53 (95% CI = [.16, .94]), *p* = .005, BF_Inclusion_ = 17.1. This result reflects a higher similarity of gaze patterns, across motivational conditions, for images that were remembered (hits), compared to images that were forgotten (misses; which is in line with findings from previous studies: e.g., Cooper et al., 2017; Foulsham & Kingstone, 2013). The Motivation x Memory Accuracy interaction was not statistically significant, *F*(2,66) = .76, *f*= .15 (95% CI = [.00, .36]), *p* = .472, BF_Exclusion_ = 2.3.

##### Encoding-regulation similarity

Encoding-regulation similarity was assessed by comparing the gaze pattern of each image in the memory-encoding phase to the gaze pattern of the subsequent blurred image in the memory-regulation phase (see solid arrow in Fig. 3A). The ANOVA revealed a significant main effect of motivation, *F*(2,66) = 10.72, *f* = .57 (95% CI = [.28, .84]), *p* < .001, BF_Inclusion_ = 43714.9. Post hoc comparisons showed that similarity scores in the remember condition were significantly higher than similarity scores in the control, *t*(33) = 3.07, *p* = .013, *d* = .53 (95% CI = [.04, 1.00]), BF_10_ = 8.9, and forget, *t*(33) = 3.81, *p* = .002, *d* = .65 (95% CI = [.16, 1.14]), BF_10_ = 51.7, conditions. These results suggest that participants looked more similarly at a blurred and its preceding scene image when motivated to remember (versus forget) the scene image. Moreover, the ANOVA revealed a significant main effect of memory accuracy, *F*(1,33) = 21.88, *f* = .81 (95% CI = [.40, 1.30]), *p* < .001, BF_Inclusion_ = 59.6, reflecting *more* similar gaze patterns, across motivational conditions, for later forgotten (misses) compared to later remembered (hits) images (see Fig. 3B). This result provides initial evidence that across-stimulus gaze similarity may interfere with subsequent memory. The Motivation x Memory Accuracy interaction was not statistically significant, *F*(2,66) = .16, *f* = .07 (95% CI = [.00, .24]), *p* = .857, BF_Exclusion_ = 2.6. Thus, while extrinsic motivation was found to influence gaze similarity between encoding and regulation phases, it did not modulate the similarity-memory association.

##### Global encoding-regulation similarity

To further asses the across-stimulus similarity and memory relationship, a more global type of similarity – similar to that used in previous neural reinstatement studies – was computed (see LaRocque et al., 2013). First, for each participant, stimuli were back-sorted according to subsequent memory (hits versus misses; see LaRocque et al., 2013; Paller & Wagner, 2002; Wagner, 1998). Then, global similarity was computed separately for hits and misses. Specifically, the gaze pattern of each hit (or miss) in the memory-encoding phase was compared to the gaze patterns of all other hits (or misses) in the memory-regulation phase (see dashed arrows in Fig. 3A). The ANOVA on this similarity measure revealed a significant main effect of memory accuracy, *F*(1,33) = 21.64, *f* = .81 (95% CI = [.40, 1.29]), *p* < .001, BF_Inclusion_ = 6.8 x 10^6^, indicating higher similarity scores for misses compared to hits (across motivational conditions; see Fig. 3B). This result supports the idea that across-stimulus gaze similarity may interfere with memory. All other effects did not reach statistical significance (*p*’s > .05). Please see the online Supplementary Information for a direct comparison of (local) encoding-regulation and global encoding-regulation similarity scores.

##### Global encoding-encoding similarity

As another global measure, we also compared the gaze pattern of each hit (or miss) in the memory-encoding phase with the gaze patterns of all other hits (or misses) in the memory-encoding phase (see dotted arrows in Fig. 3A). The ANOVA on this global similarity measure also revealed a significant main effect of memory accuracy, *F*(1,33) = 31.40, *f* = .98 (95% CI = [.53, 1.50]), *p* < .001, BF_Inclusion_ = 2.7 x 10^14^, indicating higher similarity scores for misses compared to hits (across motivational conditions; see Fig. 3B). Again, this result suggests that across-stimulus similarity may interfere with later memory. The main effect of motivation was not statistically significant and although the Motivation x Memory Accuracy effect did reach significance, *F*(2,66) = 4.13, *f* = .35 (95% CI = [.04, .60]), *p* = .020, it could not be confirmed by the BF_Inclusion_ = .2 (BF_Exclusion_ = 4.2).

In summary, the results of Experiment 1 provide preliminary evidence that: (1) when different images are encoded with more similar gaze patterns, their subsequent memory is reduced, and (2) that this similarity-memory relationship is unaffected by extrinsic motivation. For the ease of this and later discussion, we will refer to the former effect as *the across-stimulus interference effect*. It should be noted that this effect was observed both when comparing the gaze patterns during viewing of scene and other scene images (global encoding-encoding similarity) as well as when comparing the gaze patterns during viewing of scene and blurred scene images (global encoding-regulation similarity). The blurred images were however blurred versions of the encoded scene images. Hence, although they might have appeared differently, there was clearly some “perceptual overlap”. The visual distraction studies described in the introduction suggest that only when there is such perceptual overlap between stimuli, memory will be disrupted (e.g., Borst et al., 2012; Burin et al., 2007). Taking this into account, we set up Experiment 2 and changed the blurred scenes to pink noise images.

### Experiment 2

The experimental procedures of Experiment 2 were identical to Experiment 1, with the exception that blurred images were replaced with pink noise images in the memory-regulation phase (see Fig. 1). These pink noise images were randomly created and had no prior connection, nor perceptual overlap, with the encoded scene images. Although pink noise images are known to elicit an overall less explorative scanning pattern when compared to natural images, they induce a greater level of visual exploration when compared to white noise images – i.e., larger saccades, higher fixation-rate, and shorter fixation-durations (e.g., Jansen et al., 2009; Kaspar & König, 2011).

#### Recognition performance

Memory performance in the recognition test was analyzed using a repeated measures ANOVA, comparing recognition rates in the three motivational conditions: remember (*M* = 70%, 95% CI = [65, 75]), control (*M* = 60%, 95% CI = [54, 66]) and forget (*M* = 59%, 95% CI = [54, 64]). The ANOVA yielded a significant effect of motivation, *F*(2,78) = 13.60, *f* = .59 (95% CI = [.33, .85]), *p* < .001, BF_Inclusion_ = 2075.7, which was followed by post hoc comparisons. These comparisons revealed significantly higher memory in the remember condition compared to the control, *t*(39) = 4.20, *p* < .001, *d* = .66 (95% CI = [.21, 1.11]), BF_10_ = 168.6, and forget, *t*(39) = 4.85, *p* < .001, *d* = .77 (95% CI = [.31, 1.22]), BF_10_ = 1046.2, conditions. No difference in memory was found between the control and forget conditions (*p* > .05). Similar to in Experiment 1, these results suggest that participants succeeded to enhance, but not suppress, their memory.

Analyses of image memorability revealed that different participants forgot different pictures (see online Supplementary Information), as in Experiment 1. Thus, any observed differences in gaze behavior between remembered (hits) and forgotten (misses) images are unlikely to be explained by actual differences between the images themselves.

#### Skin conductance response analyses

Skin Conductance Responses (SCRs) were defined as the maximal increase in skin conductance during the 1-5 s after stimulus onset (klein Selle et al., 2016, 2017). The raw SCR data from the memory-encoding, memory-regulation and memory-test phases were analyzed separately with a Motivation (remember, control, forget) x Memory Accuracy (hits vs. misses) repeated measures ANOVA.

The ANOVA of the memory-encoding data revealed a significant main effect of memory accuracy, *F*(1,37) = 4.33, *f* = .34 (95% CI = [.00, .70]), *p* = .044, which was not confirmed by the BF_Inclusion_ = .4. Surprisingly however, SCRs during encoding were larger for subsequently forgotten (misses; *M* = .17 μS, 95% CI = [.11, .22]) compared to subsequently remembered (hits; *M* = .14 μS, 95% CI = [.09, .19]) images. Hence, differences in arousal (as measured by the SCR) during encoding are unlikely to explain later recognition-memory. All other effects were not statistically significant (all *p*’s > .05).

The ANOVA of the memory-test data also revealed a significant main effect of memory accuracy, *F*(1,38) = 8.00, *f* = .46 (95% CI = [.12, .83]), *p* = .007, BF_Inclusion_ = 10.3. This time, however, larger SCRs were observed for hits (*M* = .12 μS, 95% CI = [.08, .17]) compared to misses (*M* = .09 μS, 95% CI = [.05, .13]). In other words, memory success was reflected also in enhanced SCR amplitudes, which is consistent with the memory detection literature (klein Selle et al., 2016, 2017; Meijer et al., 2014). The ANOVA of the memory-regulation data revealed no statistically significant effects (all *p*’s > .05).

#### Gaze-similarity analyses

As for the SCR data, all gaze-similarity data were analyzed with a Motivation (remember, control, forget) x Memory Accuracy (hits vs. misses) repeated measures ANOVA. Results are depicted in Fig. 3C.

##### Encoding-test similarity

The ANOVA revealed a significant main effect of memory accuracy, *F*(1,35) = 14.05, *f* = .63 (95% CI = [.26, 1.06]), *p* < .001, BF_Inclusion_ = 8.3. This result indicates that, across motivational conditions, remembered images were scanned in a more similar manner during encoding and test, compared to forgotten images (which is in line with findings from Experiment 1 and previous studies; e.g., Cooper et al., 2017; Foulsham & Kingstone, 2013). All other effects were not statistically significant (*p*’s > .05).

##### Encoding-regulation similarity

The ANOVA revealed a significant main effect of motivation, *F*(2,68) = 5.74, *f* = .41 (95% CI = [.13, .66]), ε = .72, *p* = .011, BF_Inclusion_ = 97.6. Post hoc comparisons showed that similarity scores in the remember condition were significantly higher than similarity scores in the forget condition, *t*(35) = 2.66, *p* = .035, *d* = .46 (95% CI = [-.02, .93]), BF_10_ = 3.7; the other two comparisons did not reach statistical significance (both *p*’s > .05). These results suggest that participants looked more similarly at a pink noise image and its preceding scene image when motivated to remember (versus forget) the scene image. All other effects were not statistically significant (both *p*’s > .05). Hence, there is no support for the proposed *across-stimulus interference effect* (BF_Inclusion_ for the main effect of Memory Accuracy = .4), nor for the idea that this effect may be modulated by extrinsic motivation (BF_Inclusion_ for the Motivation x Memory Accuracy interaction = .1).

##### Global encoding-regulation similarity

The ANOVA revealed no significant effects (all *p*’s > .05). Again, this means that there is no support for the suggested *across-stimulus interference effect* (BF_Inclusion_ = 1.1). Please see the online Supplementary Information for a direct comparison of (local) encoding-regulation and global encoding-regulation similarity scores.

##### Global encoding-encoding similarity

The ANOVA revealed a trend towards significance for the main effect of memory accuracy, *F*(1,35) = 3.90, *f* = .33 (95% CI = [.00, .70]), *p* = .056, with a BF_Inclusion_ of 16.4, reflecting higher similarity scores for misses compared to hits (across motivational conditions). As in Experiment 1, this result supports the idea that across-stimulus similarity interferes with later memory. All other effects were not statistically significant (all *p*’s > .05).

Taken together, the results of Experiment 2 are partly consistent with those reported in Experiment 1. Specifically, we found strong (Bayesian) evidence for the suggested *across-stimulus interference effect*, for 1 out of 3 comparison types. Specifically, when gaze-scanning patterns were compared across scenes and other scene images (during encoding), more similar explorations were related to lower subsequent memory. On the other hand, when gaze-scanning patterns were compared across scenes and pink noise images, exploration similarity seemed extraneous to future memory success. These results provide some support for the hypothesis that there needs to be a certain degree of perceptual (information) overlap between the encoded images – as was the case for the scene and blurred scene images in Experiment 1, but not for the scene and pink noise images in Experiment 2 – for gaze similarity to affect later memory. In addition, as found in Experiment 1, although extrinsic motivation affected the way in which participants viewed the pink noise images (i.e., encoding-regulation similarity was highest when motivated to remember), it did not mediate the observed similarity-memory relationship.

## Discussion

How we look out at the world influences the way in which we remember it. Until now, studies on gaze-pattern similarity have focused on how we look at identical stimuli (e.g., images) during encoding and memory testing (e.g., Cooper et al., 2017; Foulsham & Kingstone, 2013). These studies showed that greater encoding-test gaze similarity relates to better memory (as replicated here). The present study extends these findings by examining how we look at various different stimuli during encoding and whether this affects future memory success. Based on existing neural reinstatement work, we hypothesized that high levels of across-stimulus gaze similarity would relate to low levels of memory. Furthermore, we examined whether the motivation to either forget or remember affects the hypothesized similarity-memory relationship.

When considering the first aim of this study, our results provide initial evidence that higher levels of gaze similarity (during encoding) do indeed relate to lower levels of subsequent memory. Experiment 2, however, suggests that there may be a limit to this *across-stimulus interference effect* and that it depends on the perceptual similarity (overlap) between the stimuli – namely, there needs to be some perceptual overlap between the relevant stimuli (as was true for the scene-blurred scene, but not for the scene-pink noise comparisons) for similar gaze patterns to disrupt memory. Although this idea fits with earlier findings from the visual distraction literature (e.g., Borst et al., 2012; Burin et al., 2007) – i.e., only similar distracters disrupt memory – it remains to be determined whether the observed memory disruption in these studies might be driven by underlying gaze-similarity effects (none of these previous studies used eye tracking).

As the suggested *across-stimulus interference effect* seems to depend on the perceptual overlap between stimuli, one may wonder what happens when eye-movements are reinstated in the absence of a stimulus (i.e., “looking-at-nothing”). Previous studies examined exactly this and showed that more similar exploration patterns during encoding and delay may actually benefit memory (e.g., Laeng et al., 2014; Olsen et al., 2014; Wynn et al., 2018). Interestingly, this suggests a non-linear effect of gaze reinstatement, with beneficial memory effects when no other stimulus is presented, no memory effects when the encoded stimuli are perceptually different, and disruptive memory effects when the encoded stimuli are perceptually similar. This could potentially provide an intriguing avenue for future research.

Integrating the present findings with the neuroimaging literature on pattern similarity reveals an interesting commonality. Just as observed for gaze similarity here, activation similarity (i.e., reduced pattern separation) in the hippocampus (but not in other medial temporal lobe regions) has been found to interfere with memory (e.g., Chanales et al., 2017; Favila et al., 2016; Hulbert & Norman, 2015; LaRocque et al., 2013). This may however not be that surprising, considering that the hippocampus and the oculomotor system are anatomically well connected through an extensive set of polysynaptic pathways (Shen et al., 2016). In line with this reasoning, individuals with amnesia whose damage includes the hippocampus, show alterations in their gaze patterns (Olsen et al., 2015; see also Voss et al., 2011). In addition, there is recent evidence that visual sampling during encoding can predict hippocampal activity in neurologically intact individuals (Liu et al., 2017; Ringo et al., 1994; Ryals et al., 2015). Finally, hippocampus activation was shown to be linked to the expression of relational memory in viewing patterns even when explicit retrieval failed (Hannula & Ranganath, 2009). Taken together, without assuming causality, these findings suggest that the hippocampal and oculomotor networks are inherently linked and raise the possibility that the observed gaze similarity effects, in the present and previous studies, may be related to activation similarity in the hippocampus (see also Bone et al., 2018).

An alternative explanation of the observed *across-stimulus interference effect* could be that the forgotten images were simply less perceptually engaging, less distinctive and less memorable, consequently inducing more similar gaze patterns. Two of our findings however argue against this explanation. First, our SCR analysis in Experiment 2 revealed that the forgotten images (i.e., misses) induced larger SCRs than the remembered images (i.e., hits) during encoding. This suggests that the forgotten images were encoded with a higher level of arousal, which would be unlikely if these images were less engaging and less memorable. Second, different participants forgot different pictures, suggesting that there were no consistent differences in the distinctiveness and memorability of the images.

If the *across-stimulus interference effect* proves to be stable in future replications (preferably with different types of stimuli), a potential mechanism for successful memory regulation might have been illuminated. A deeper understanding of such mechanism may hold clinical implications for the treatment of different psychopathologies characterized by the intrusion of unwanted memories: e.g., posttraumatic-stress disorder, obsessive-compulsive disorder, and depression. Intrusions are typically vivid, detailed, unexpected and uncontrollable. To resist such intrusions, people often attempt to self-distract and avoid triggers, which paradoxically increase thought frequency, hyper-vigilance, and negative appraisal of the intrusions (Purdon, 2004). If, however, there is a causal connection between gaze reinstatement and successful memory suppression, it could pave the way for alternative clinical interventions. These could for instance involve simple experimental manipulations of visual exploration patterns to manipulate gaze reinstatement. The ease by which eye movements can be measured, and controlled, presents an important advantage over the more covert neural measures. Nevertheless, considering again the close functional and anatomical connection between the visual and hippocampal systems (Bone et al., 2018; Bosch et al., 2014; Liu et al., 2017; Pacheco Estefan et al., 2019; Ringo et al., 1994; Ryals et al., 2015; Shen et al., 2016), gaze manipulation may possibly also affect the underlying neural processes.

Regarding the second aim of this study, our results suggest that the motivation to either forget or remember does not mediate the observed gaze-similarity and memory relationship. Motivated memory did however affect (encoding-regulation) similarity. Specifically, participants looked more similarly at either a blurred or pink noise image and a preceding scene image, when motivated to remember the scene image. Interestingly, only 2-3 participants, in each experiment, reported to have purposely changed their gaze behavior depending on their motivational state. This suggests that participants were unconsciously reinstating their gaze when trying to hold on to (and remember) an image. At any case, whether conscious or not, this gaze-reinstatement effect did not benefit memory, since across motivational conditions, higher encoding-regulation similarity scores were observed for misses compared to hits.

The absence of a memory suppression effect (memory performance in forget condition < memory performance in control condition) follows earlier inconsistent findings in the motivated memory literature. When considering previous research that used a directed forgetting paradigm (like in the present study), initial studies did not include a control condition and only compared memory in a remember condition with memory in a forget condition (e.g., Fawcett & Taylor, 2008; Wylie et al., 2008). Clearly, if a difference is observed, this difference could have resulted from both a memory facilitation (remember > control) and a memory suppression (forget < control) effect. More recent studies have included a control condition and found evidence solely for memory facilitation (as in the present study; e.g., Gao et al., 2016; Zwissler et al., 2015). When considering studies that used a think-no-think paradigm, the evidence for memory suppression seems stronger (e.g., Anderson & Green, 2001; Depue et al., 2007, 2016), however contradictory evidence exists (e.g., Bulevich et al., 2006; Mecklinger et al., 2009). In a typical think-no-think study, participants are presented with item-pairs during encoding, whereas during testing they are presented with only one item of a pair. Participants are then asked to try and think or not to think of the paired associate. Thus, while actual item-memories are suppressed in the directed forgetting paradigm, associative memories are suppressed in the think-no-think paradigm. This difference in the type of suppression could possibly explain the discrepancy in findings with the two paradigms (see also Rosenfeld et al., 2017).

One limitation of the present study which should be acknowledged is that, in certain participants, the blurred and pink noise images induced very little visual exploration. Hence, after excluding all trials with less than 3 fixations, the eye-tracking data of some participants had to be excluded. As mentioned earlier, it is common that pink noise images (and even more so white noise images) induce more fixations to the center of the screen than natural images (e.g., Jansen et al., 2009; Kaspar & König, 2011). Hence, future studies should consider using other types of more complex stimuli (e.g., fractals), in order to stimulate eye movements and reduce the central fixation bias.

Taken together, the present study provides preliminary evidence for the idea that across-stimulus gaze similarity during encoding interferes with subsequent memory. In other words, when different stimuli are encoded with more similar gaze-scanning patterns, later memory reports are hampered. In addition, while this effect seems dependent upon the amount of perceptual overlap between the encoded stimuli, it seems unaffected by extrinsic motivation to either forget or remember. Although these findings translate to established neural reinstatement effects, the implications are divergent. Specifically, the ability to control eye movements raises the possibility that direct manipulations of gaze similarity (during encoding) would affect later recognition. If so, it could potentially serve as a mechanism to aid the control of unwanted memories.

## Methods

### Participants

A total of seventy-nine participants of the Hebrew University of Jerusalem (HUJI) took part in the study. Thirty-six undergraduate students (30 women), with an age range of 20-35 years (*M* = 23.9 years, *SD* = 3.4 years), participated in Experiment 1. Forty-three undergraduate students (25 women), with an age range of 18-31 years (*M* = 24.0 years, *SD* = 2.7 years), participated in Experiment 2. All participants were native speakers of Hebrew and received either course credits or an average payment of 55 new Israeli shekels (NIS; ~16 USD) for their participation. The experiments were approved by the ethical committee of the Faculty of Social Sciences of the HUJI and were performed in accordance with the relevant guidelines and regulations. Each participant read and signed an informed consent form indicating that participation is voluntary and could be stopped at any stage.

### Stimuli

A total of 180 color scene images were chosen from two different databases: The INRIA Holidays dataset (Jegou et al., 2008) and the Nencki Affective Picture System (Marchewka et al., 2014); all chosen images were either natural or urban scene images. The 180 images were randomly divided into two sets of 90 images, with an equal number of natural and urban scenes within each set. Importantly, during the memory-encoding phase (of session 1), only one picture-set was presented to participants (set 1 for uneven participant numbers, set 2 for even participant numbers), whereas in the memory-test phase (of session 2), both picture-sets were presented to participants (see Procedure below).

During the memory-regulation phase (of session 1), participants were presented either with blurred, black-and-white, versions of the encoded scene images (Experiment 1) or pink noise images (Experiment 2). Image blurring was accomplished using a free online blurring tool (https://pinetools.com/blur-image; stack blur, 100 radius). Pink noise images (90 in total) were created using a Matlab utility function (the function can be downloaded from: https://github.com/kendrickkay/knkutils). Please note that while the presentation of the blurred scenes depended on the preceding scene stimulus (Experiment 1), the presentation of pink noise images was randomly determined (Experiment 2). All images were presented on a Syncmaster monitor at a resolution of 1024 x 768 pixels (screen resolution was 1920 x 1080), at a viewing distance of approximately 60 cm.

### Procedure

All participants underwent two testing sessions which were separated by 2-3 days (see Fig. 1). In both sessions, a short break was inserted halfway (splitting each session into two blocks), to enable a short rest for the participants and keep their vigilance. The sessions were structured as follows:

#### Session 1

Once informed consent was obtained, the experimenter identified the participant’s dominant eye and attached the electrodes for measuring electrodermal activity. Next, skin conductance was measured during 1 minute of rest (i.e., baseline period). When ready, participants were instructed about the exact experimental procedures: The experiment consisted of 90 trials, and each trial began with the presentation of 1 out of 90 colored scene-pictures for 3000 ms (i.e., memory-encoding phase). Immediately after each display, participants were presented with both an auditory message and visual cue (for 650 ms) instructing them to: either do nothing (white fixation dot), forget (purple fixation dot) or remember (green fixation dot) the previously presented picture. The cue condition of each picture was randomly determined and no more than 2 identical cues appeared consecutively. Following the cue, participants were presented either with a blurred version of the previously presented scene (Experiment 1) or a pink noise image (Experiment 2) for 3000 ms (i.e., memory-regulation phase). In order to enhance motivation, participants were asked to imagine that they are guilty of a crime and that all pictures followed by a forget cue are related to the crime, whereas all pictures followed by a remember cue are related to their alibi. Moreover, participants were told that in the next experimental session (2-3 days later), they would undergo a polygraph test in which they can win a 15 NIS (~4.3 USD) bonus if the polygraph test: (1) will not connect them to the crime-related pictures, but (2) will connect them to the alibi-related pictures. After the experimenter ensured that the instructions were understood, the eye-tracker was set up and a standard nine-point calibration and validation procedure (Experiment Builder SR research; Ontario Canada) was performed. Finally, before starting the actual experiment, all participants also underwent a short practice phase (of 3 trials) to familiarize them with the procedure.

#### Session 2

After the experimenter attached the skin conductance electrodes, participants were explained that they would undergo a regular recognition-memory test, not a polygraph test (as told in session 1). In this memory test, participants were presented with the 90 studied pictures from session 1 of the experiment and 90 foils taken from the unstudied picture set (each picture was presented for 3000 ms; i.e., memory-test phase). After the picture disappeared, participants were asked whether or not it had been presented during session 1. Importantly, they were told to disregard the previous forget and remember instructions, and all earlier presented stimuli, regardless of previous instructions, should be endorsed with yes. Participants were given unlimited time to make their responses and accuracy was encouraged by promising a monetary bonus (i.e., 15 NIS) if at least 75% of their answers were correct (i.e., “yes” responses to studied items, “no” responses to foils). After the experimenter ensured that all experimental procedures were understood, the eye-tracker was set up and a standard nine-point calibration and validation procedure was performed. When the participant reported to be ready, the experimental session started.

At the end of session 2, participants received a paper-and-pencil questionnaire in which they were asked to rate, on a scale 1 to 6 (1 = *not at all*, 6 = *very much*), their overall motivation to remember versus forget the images (that were followed by remember versus forget cues, respectively), as well as their efforts to remember versus forget the images (that were followed by remember versus forget cues, respectively). Furthermore, participants were asked to verbally describe any strategies used to help them remember or forget. Analyses of the questionnaire data are presented in the online Supplementary Information. Finally, all participants were debriefed and compensated for their participation in the experiment.

### Data acquisition and reduction

The experiment was conducted in a sound attenuated room with dedicated airconditioning in order to keep the temperature stable. The apparatus included a Biopac MP160 system (BIOPAC Systems, Inc., Camino Goleta, CA) to measure skin conductance and a ThinkCentre M Series computer to save the relevant data. Skin conductance was obtained with a sampling rate of 1000 Hz and two Ag/AgCl electrodes (1.6-cm diameter) which were placed on the distal phalanges of the left index and left ring finger. Due to technical issues in Experiment 1, SCRs were analyzed only in Experiment 2.

An EyeLink 1000-plus table-mount setup was used to measure the eye-movement data and a SilverStone computer saved the relevant data. Eye-movement data were parsed into saccades and fixations using EyeLink’s standard parser configuration: samples were defined as a saccade when the deviation of consecutive samples exceeded 30°/s velocity or 8000°/s^2^ acceleration. Samples gathered from time intervals between saccades were defined as fixations.

In Experiment 1, after disqualifying trials with less than 3 fixations (2.3% of all trials), all eye-tracking data of two (out of thirty-six) participants were excluded from analysis because more than 20% of their data (from the memory-regulation phase) were removed. Thus, all eye-movement analyses of Experiment 1 were based on data of 34 participants. An a-priori power analysis revealed that this sample size allows for detecting a medium effect size (i.e., Cohen’s *d* of 0.50) with a statistical power of at least 0.80.

In Experiment 2, after disqualifying trials with less than 3 fixations (2.3% of all trials), all eye-tracking data of three (out of forty-three) participants were excluded from analysis because more than 20% of their data (from the memory-regulation phase) were removed. In addition, depending on the type of similarity scores analyzed (i.e., encoding-test, encoding-regulation, global encoding-regulation, global encoding-encoding), the eye-tracking data of either one or two additional participants were disqualified because no “misses” data remained in one of the experimental conditions. Finally, all data of three participants were disqualified either because they did not show up to the second part of the experiment or because of noncompliance. Thus, all eye-movement analyses were based on data of 35-36 participants.

In Experiment 2, we also analyzed SCRs that were defined as the maximal increase in skin conductance during the 1-5 s after stimulus onset (klein Selle et al., 2016, 2017). Although raw SCRs were analyzed, all SCR values were standardized to remove outliers (i.e., standard score is larger than 5 or smaller than −5) as well as trials with excessive movements (i.e., standard score is larger than 0 when a movement occurred). A total of 1.5% of SCRs were eliminated from the memory-encoding phase, a total of 1.5% of SCRs were eliminated from the memory-regulation phase, and a total of 1.1% of SCRs were removed from the memory-test phase. Importantly, the within-subject standardization was performed within experimental blocks (before and after the break), minimizing habituation effects (see Ben-Shakhar & Elaad, 2002; Elaad & Ben-Shakhar, 1997).

In addition, skin-conductance nonresponsivity was determined after the elimination of single items (similar to klein Selle et al., 2016, 2017). Specifically, participants in which the standard deviation across trials in both the first and second block of a session was below 0.01 μS, were considered to be nonresponders and their SCR data were eliminated from all analyses. In case of nonresponsivity in either the first or the second block, only the data from the respective trials were removed^3^. For the memory-encoding phase, this led to the removal of all SCR data of 1 participant, the SCR data of the first block of 2 participants, as well as the SCR data of the second block of another 2 participants. For the memory-regulation phase, this led to the removal of the SCR data from the first block of 2 participants and the SCR data from the second block of another 3 participants. For the memory-test phase, this led to the removal of the SCR data of the second block of 2 participants. Finally, the skin conductance data of one additional participant were disqualified because no “misses” data remained in one of the experimental conditions. Thus, all skin conductance analyses were based on data of either 38 or 39 participants.

### Data analyses

Gaze similarity was computed using the ScanMatch toolbox for Matlab (The MathWorks, Natick, MA; Cristino et al., 2010). Using this method, each fixation sequence is spatially and temporally binned and then recoded to create a sequence of letters that retains fixation location, order, and time information. Pairs of these sequences are then compared using the Needleman-Wunsch algorithm (borrowed from the field of genetics) to find the optimal alignment between a pair. The correspondence between two sequences is expressed by a normalized similarity score (0 = *no correspondence*, 1 = *identical*; see Fig. 2) – which is inversely related to the number of actions needed to transform one sequence into the other. In the present study, ScanMatch was run using a 12 x 8 bin ROI grid (the default). Furthermore, for temporal binning we applied a value of 50 ms, which has been demonstrated to give the most accurate sampling across a wide variety of fixation durations (see Cristino et al., 2010). Finally, a substitution matrix threshold of 4 was used, which was 2 times the standard deviation of the ‘gridded’ saccade size (i.e., threshold = 2 x standard deviation (mean saccadic amplitude in pixels) / (Xres / Xbin); with Xres = X resolution of the stimuli, and Xbin = number of bins horizontally). This means that the alignment algorithm aimed to align only regions that were maximum 4 bins apart.

Results were analyzed using Matlab R2016a (The MathWorks, Natick, MA) and R software (version 3.6.1). Mean recognition, SCR and gaze-similarity scores were subjected to repeated-measures ANOVAs; for ANOVAs involving more than one degree of freedom in the enumerator, the Greenhouse-Geisser procedure was applied when the assumption of sphericity was violated. For all post hoc comparisons, the Bonferroni-corrected *p*-value is reported. Both Cohen’s *f* and Cohen’s *d* values were computed as effect size estimates (Cohen, 1988). In addition to frequentist statistical inference, we relied on Bayesian analyses and computed Jeffreys-Zellener-Siow (JZS) Bayes factors (BFs). Please note that the default prior settings (used by R) were left unchanged. For all *t*-tests (two-sided), either the BF_10_ (quantifying the evidence favoring the alternative hypothesis) or the BF_01_ (quantifying the evidence favoring the null hypothesis) is reported. For all ANOVA main and interaction effects, either the BF_Inclusion_ or BF_Exclusion_ is reported, reflecting a comparison of all models including (or excluding) a particular effect to those without (or with) the effect. In other words, the BF_Inclusion_ can be interpreted as the evidence in the data for including an effect or interaction, similar to BF_10_ in the case of simple comparisons (see also van den Bergh et al., 2020). Therefore, the conventions used to interpret substantial/moderate support for either the null or alternative hypothesis (BF_10_ >= 3; Jeffreys, 1998) may apply also to BF_Inclusion_.

### Preregistration and data availability

All analyses (as well as the experimental design and hypotheses) of Experiment 2 were preregistered on: https://aspredicted.org/k8re8.pdf. The original data and analysis files of both Experiments 1 and 2 can be accessed on: https://osf.io/gh7ba/.

## Data Availability

Data of all participants in the two experiments are publicly available on the OSF (https://osf.io/gh7ba/).

## Code Availability

Custom code that supports the findings of this study is publicly available on the OSF (https://osf.io/gh7ba/).

## Acknowledgements

This research was funded by a grant, No. 1747/14, from the Israel Science Foundation to YP. The funders had no role in the conceptualization, design, data collection, analysis, decision to publish, or preparation of the manuscript. We wish to thank Hagit Tanami and Danna Waxman for their assistance in data collection.

## Author Contributions

NKS, MG and YP designed the study, NKS acquired and analyzed the data, MG and YP supervised the analysis, NKS wrote the manuscript, and all authors contributed to revisions of the manuscript.

## Footnotes

1 The skin conductance data of Experiment 1 was not analyzed due to technical problems. These problems were solved before starting Experiment 2.

2 In addition to the image-memorability analyses, the online Supplementary Information also includes: analyses of reaction times during the recognition test, subjective motivation and effort ratings, potential strategies for remembering or forgetting the scenes, respectively, and local versus global gaze similarity analyses.

3 Please note we did not pre-register the removal of SCR data based on nonresponsivity. Nonetheless, when including the ‘nonresponding’ blocks, similar results are observed.

