## Supplementary Information for "When Gaze-Pattern Similarity May Interfere With Future Memory"

### Experiment 1

#### Subjective ratings

The paper-and-pencil questionnaire data acquired at the end of session 2 were analyzed using paired samples *t*-tests. These tests revealed a significant difference between the self-reported motivation to remember ($M$ = 5.33, 95% CI = [5.03, 5.63]) and forget ($M$ = 4.53, 95% CI = [4.15, 4.91]), $t(35) = 4.54$, $p < .001$, $d = .76$ (95% CI = [.38, 1.12]), $BF_{10}$ = 376.9. Similar results were obtained for self-reported efforts to remember ($M$ = 5.22, 95% CI = [4.87, 5.57]) and forget ($M$ = 4.31, 95% CI = [3.92, 4.69]), $t(35) = 4.11$, $p < .001$, $d = .69$ (95% CI = [.32, 1.04]), $BF_{10}$ = 118.1. Taken together, participants indicated to have been more motivated, and to put more effort into their task, when instructed to remember than when instructed to forget. The most commonly reported strategies in the remember condition included: focus on specific details, creating a story around all the to-be-remembered pictures, associate the pictures with personal memories. In the forget condition, on the other hand, the most commonly reported strategies were: to think of something else, to think of the last picture that had to be remembered, to distract oneself. Only three participants indicated to have changed the way they scanned the blurred images, depending on whether they tried to forget or remember.

#### Image-memorability analyses

In order to validate that there were no inherent and consistent differences in the memorability of the experimental images (i.e., that participants did not forget the same pictures), we correlated the memory scores (1 = *hit*, 0 = *miss*) of all images (Boolean vector of 90 elements) across all possible pairs of participants. The mean correlation was low and not significantly different from zero ($r = .05$, $p = .474$). Thus, participants seemed to forget different pictures, suggesting that there were no inherent differences in the memorability of our images. Consequently, any observed differences in gaze behavior between remembered (hits) and forgotten (misses) images are unlikely to be explained by actual differences between the images themselves.

**Reaction-time analyses**

The reaction-time data (from the memory-test phase) were analyzed with a 3 x 2 repeated measures ANOVA, with motivation (remember, control, forget) and memory accuracy (hits vs. misses) as within-subject factors. The ANOVA revealed a significant main effect of memory accuracy, $F(1,35) = 4.37$, $f = .35$ (95% CI = [.00, .73]), $p = .044$, which could not be confirmed by the $BF_{Inclusion} = .6$. This result reflects the higher mean reaction time for misses ($M = 684$ ms, 95% CI = [611, 757]) compared to hits ($M = 610$ ms, 95% CI = [537, 683]), across motivational conditions. All other effects were not statistically significant ($p$'s > .05).

**Local versus global encoding-regulation similarity**

In order to examine the specificity of the (local) encoding-regulation and global encoding-regulation effects, we submitted the respective similarity scores to a repeated measures ANOVA, with motivation (remember, control, forget), memory accuracy (hits vs. misses), and similarity-type (local vs. global) as within-subject factors. The ANOVA revealed a significant main effect of similarity-type, $F(1,33) = 119.53$, $f = 1.90$ (95% CI = [1.24, 2.73]), $p < .001$, $BF_{Inclusion} > 10^{12}$, reflecting higher similarity scores for local (scene versus subsequent blurred image) than global (scene versus all other blurred images) comparisons. Moreover, the ANOVA revealed a significant main effect of motivation, $F(2,66) = 6.53$, $f = .45$ (95% CI = [.16, .70]), $p = .003$, $BF_{Inclusion} = 6.1 \times 10^9$, as well as a significant Motivation x Similarity-Type interaction, $F(2,66) = 13.86$, $f = .65$ (95% CI = [.35, .93]), $\varepsilon = .85$, $p < .001$, $BF_{Inclusion} = 2.1 \times 10^9$. The interaction reflects the fact that the main effect of motivation was significant for local, but not for global, encoding-regulation similarity scores (please see Results section). In addition, the ANOVA revealed a significant main effect of memory accuracy, $F(1,33) = 28.43$, $f = .93$ (95% CI = [.49, 1.44]), $p < .001$, $BF_{Inclusion} = 972.5$, reflecting higher similarity scores for misses compared to hits (across motivational conditions and similarity-types). This result supports the idea that across-stimulus similarity may interfere with later memory. No other main or interaction effects were statistically significant (all *p*'s > .05). Altogether these results substantiate our main findings.

**Experiment 2**

### Subjective ratings

The paper-and-pencil questionnaire data were analyzed similarly as in Experiment 1 using paired samples *t*-tests. These tests revealed a significant difference between the self-reported motivation to remember ($M = 5.49$, 95% CI = [5.24, 5.74]) and forget ($M = 4.76$, 95% CI = [4.42, 5.09]), $t(40) = 3.89$, $p < .001$, $d = .61$ (95% CI = [.27, .94]), $BF_{10} = 73.8$. Similar results were obtained for self-reported efforts to remember ($M = 5.32$, 95% CI = [5.05, 5.59]) and forget ($M = 4.02$, 95% CI = [3.54, 4.51]), $t(40) = 5.22$, $p < .001$, $d = .82$ (95% CI = [.46, 1.17]), $BF_{10} = 3299.8$. Taken together, participants indicated to have been more motivated, and to put more effort into their task, when instructed to remember than when instructed to forget, which is consistent with the results of Experiment 1. The most commonly named strategies in the remember condition included: naming the pictures, creating a story around all to-be-remembered pictures, associate the pictures with personal memories. In the forget condition, on the other hand, the most commonly named strategies included: to think of something else, to think of the last picture that had to be remembered, to distract oneself. Only two participants indicated to have changed the way they scanned the pink noise images, depending on whether they tried to forget or remember.

**Image-memorability analyses**

In order to validate that there were no inherent differences in the memorability of the experimental images (i.e., that participants did not forget the same pictures), we correlated the 90 memory scores (1 = *hit*, 0 = *miss)* for each of the images, across all possible pairs of participants. As in Experiment 1, the mean correlation was low and not significantly different from zero ($r = .07$, $p = .449$).

**Reaction-time analyses**

The reaction-time data (from the memory-test phase) were analyzed with a 3 x 2 repeated measures ANOVA, with motivation (remember, control, forget) and memory accuracy (hits vs. misses) as within-subject factors. The ANOVA revealed a significant main effect of memory accuracy, $F(1,38) = 7.57$, $f = .45$ (95% CI = [.11, .82]), $p = .009$, $BF_{Inclusion}$ = 14.2. This result reflects the higher mean reaction time for misses ($M = 796$ ms, 95% CI = [673, 919]) compared to hits ($M = 672$ ms, 95% CI = [549, 795]), across motivational conditions. All other effects did not reach statistical significance ($p$'s > .05).

**Local versus global encoding-regulation similarity**

In order to examine the specificity of the (local) encoding-regulation and global encoding-regulation effects, we submitted the respective similarity scores to a repeated measures ANOVA, with motivation (remember, control, forget), memory accuracy (hits vs. misses), and similarity-type (local vs. global) as within-subject factors. The ANOVA revealed a significant main effect of similarity-type, $F(1,34) = 19.05$, $f = .75$ (95% CI = [.35, 1.21]), $p < .001$, $BF_{Inclusion}$ = 7.2, reflecting higher similarity scores for local (scene versus subsequent pink noise image) than global (scene versus all other pink noise images) comparisons. Moreover, the ANOVA revealed a significant main effect of motivation, $F(2,68) = 5.90$, $f = .42$ (95% CI = [.13, .67]), $p = .004$, $BF_{Inclusion}$ = 412.8, as well as a significant Motivation x Similarity-Type interaction, $F(2,68) = 4.46$, $f = .36$ (95% CI = [.07, .60]), $\varepsilon = .65$, $p = .031$,

$BF_{Inclusion}$ = 14.9. The interaction reflects the fact that the main effect of motivation was significant for local, but not for global, encoding-regulation similarity scores (please see Results section). No other main or interaction effects were statistically significant (all *p*'s > .05). Thus, there was also no support for the suggested across-stimulus interference effect ($BF_{Inclusion}$ for the main effect of Memory Accuracy = .7). Altogether these results substantiate our main findings.
